# Differential CTCF binding in motor neurons and lymphocytes

**DOI:** 10.1101/2021.03.19.435989

**Authors:** Belkis Atasever Arslan, Scott T Brady, Gamze Gunal Sadik

## Abstract

Transcriptional regulation of protein-coding genes is a primary control mechanism of cellular function. Similarities in regulation of expression for select genes between lymphocytes and neurons have led to proposals that such genes may be useful biomarkers for some neurological disorders that can be monitored via patient lymphocyte populations. Examination of shared molecular mechanisms underlying neurogenesis and lymphocyte differentiation may give help to identify relevant pathways and suggest additional biomarkers in lymphocytes that are relevant to neurological disorders. In this study, we analysed similarities and conserved regions in several genes regulated by CCCTC-binding factor (CTCF) during lymphocyte and neuronal developmental stages. We performed epigenetic analysis of CTCF binding Trak1, Gabpa, Gabpb1, Gabpb2, Gfi1, Gfi1b gene loci at T and B lymphocytes at different developmental stages, as well as in neural progenitor cells and motor neurons. Common and shared CTCF binding events at Trak1 gene suggest additional transcriptional regulatory factors that control Trak1 gene expression levels differ in neurons and lymphocytes. Gabpb1 gene includes a common CTCF binding site shared with neurons and lymphocytes. Correlation of CTCF binding analysis and gene expression profile suggests that CTCF binding plays a role in epigenetic regulation of Gabpb1 gene. In contrast, while Gfi1a gene is phylogenetically well-conserved and CTCF sites are occupied in lymphocytes, there are no CTCF binding occupied in neurons and neural progenitor cells. Low expression levels of Gfi1s in neurons indicate that regulation of this gene is CTCF-independent in neurons. Although Gfi1b is highly homologous to Gfi1, differences in expression levels suggest that Gfi1b is critical for both lymphogenesis and neurogenesis. Neurons and lymphocytes have multiple common CTCF binding sites in the Gfi1b gene, although lineage specific transcriptional regulators that play a role in their different expression levels still need to be identified. The partial overlap in CTCF regulatory sites for some genes in neurons and lymphocytes suggest that there may be markers that can exhibit parallel changes in these cells and serve as biomarkers.

## INTRODUCTION

Transcriptional regulation of protein-coding genes is the primary control mechanism of cellular functions. Cell type diversity is critical to lymphocyte roles in the immune system as well as to neurons that form complex network structures in brain. However, mechanisms underlying this diversity remain to be elucidated. Nevertheless, similarities between lymphocytes and neurons in regulation of expression for select genes have been proposed as biomarkers for several neurological disorders that can be monitored via patient lymphocyte populations. Examination of shared molecular mechanisms underlying neurogenesis and lymphocyte differentiation may give rise to a clearer definition of relevant pathways and suggest additional biomarkers in lymphocytes relevant to neurological disorders. Identification of new targets for diagnosis and treatment of neurodegenerative disorders is crucial, as many of these disorders cannot be reliably diagnosed. In this study, we analysed similarities and conserved regions in several genes regulated by CCCTC-binding factor (CTCF) during lymphocyte and neuronal developmental stages.

CTCF is a ubiquitously expressed zinc finger protein best known for its function in transcriptional insulation (Phillips and Corces, 2009). CTCF binds to insulator regions and blocks enhancer-promoter interactions physically (Holwerda and de Laat, 2013). Thus, it can be involved in either activation or repression of gene expression. Together with cohesin, CTCF is responsible for the regulation of higher order chromatin structure, allowing and/or blocking transcriptional machinery, gene rearrangements and enhancer-promoter activity (Lee and Iyer). CTCF is well known for its functions in epigenetic regulation during lymphocyte differentiation (Gunal Sadik et. al., 2014), but has also been shown to play a role in regulation of proliferation-differentiation equilibrium of neuroprogenitor cells during cortex formation (Watson et al., 2014). As a result, examining the role of CTCF in regulating expression of specific genes in differentiation of neurons and lymphocytes may be informative.

Recent studies revealed that the co-localization of cohesin and CTCF at the variable Igh segments affects the usage of V gene segments during V(D)J recombination in pre-B cells. Cohesin and CTCF both facilitate long-range interactions between the V genes, the Eμ enhancer and some 3’ cohesin/CTCF binding sites (Gunal Sadik et. al., 2014; Guo et al., 2011; Seitan et al., 2012). CTCF is a key molecule playing critical roles in conformational changes in chromatin that determine cellular differentiation. The roles of CTCF in neurons have yet to be fully elucidated, but CTCF knock-out mice revealed significant changes in 390 transcripts of cortex and hippocampus; suggesting that it may play a role in regulating neural development (Hirayama et al., 2012). To further investigate potential roles of CTCF in neurons, we analysed shared CTCF binding sites at several genes critical for neuronal and lymphocyte development using publicly available genomics data. We determined common as well as non-shared CTCF binding sites at these genes through developmental stages of neurons and lymphocytes. We also analysed changes in expression levels of these genes in neurons and lymphocytes and determined sequence conservancy via phylogenetic analysis.

During brain development, changing local energy needs should be met during differentiation; thus, mitochondrial transport is critical for brain development (Ogawa et al., 2013). These energy needs are fulfilled by mitochondria that facilitate ATP synthesis. Proper differentiation of cells depends on availability of required energy at times of necessity. In cells like neurons that show high levels of polarization, axonal transport mechanisms are dependent on proper mitochondrial function. One protein implicated in the transport of mitochondria in axons is TRAK1, which is thought to play a critical role in the long distance trafficking of mitochondria in axons (melko and abdu) (Ogawa et al., 2013). However, TRAK1 is also expressed in lymphocytes, where distances traveled are shorter and the degree of polarization is less. Can we gain insight into TRAK1 functions shared between neurons and lymphocytes by looking at expression during differentiation of neurons and lymphocyte lineages?

Another transcription factor implicated in mitochondrial function is the GA binding protein (GABP) transcription factor, which is implicated in mitochondrial biogenesis and expression of mitochondrial genes needed for ATP production (Yang et. AL, 2014). This transcription factor binds to DNA sequences that are rich in the nucleotides guanine and adenine, and is composed of two subunits; GABPalpha and GABPbeta. GABPalpha comprises an Ets domain that binds to DNA. GABPbeta also contains four ankyrin repeats that mediate protein-protein interactions. GABPalpha/GABPbeta dimer contains a nuclear localisation signal for nuclear import. GABP regulates Pax5 gene expression and plays role in B cell differentiation (Jing et al., 2008), but little is known about its role in neuronal development.

Growth factor independence-1 (Gfi1), is a transcription factor that is involved in both neuronal and hematopoietic cell development. Deletion of Gfi1 in mice give rise to defects in sensory neurons and developmental disorders in neuroendocrine cells and causes defective hematopoietic stem cells and neutrophil deficiency. Gfi1 mediates distinct stages of lymphocyte development due to its transcriptional repressor function and it is critical for early development of T lymphocytes (Kazanjian et al., 2006). Double positive T lymphocytes arise from DN T cells that rearrange T cell receptor β-chain through expansion and differentiation. Early T cell development is blocked by Gfi1 over expression that give rise to defects in selection of productive TCR-expressing cells (Kazanjian et al., 2006). The role of Gfi1 in neuronal development is less will defined, but neurodegeneration due to mutations in Ataxin-1 (SpinoCerebellar Ataxia type 1, SCA1) is thought to be mediated by interaction with Gfi1 and loss of Gfi1 mimics the SCA1 phenotype (Tsuda et. al., 2005).

Gfi1B is 97 % homologous to Gfi1. Gfi1b is fundamental for megakaryocytic and erythroid development (van der Meer et al., 2010). Both are expressed in T-cell precursors as well (Kazanjian et al., 2006). Gfi1 is a target for p53 and regulation of hematopoietic stem cells via Gfi1 is mediated by p53. It is also involved in B cell differentaion, as Gfi1 expression decreases in mature B cells (van der Meer et al., 2010). Gfi1B is more widely expressed in adult that Gfi1, but little is known about Gfi1B function in neurons and deletion of Gif1B is embryonic lethal (Wallis et., al., 2003). However, members of this family play an important role in neuronal development/maintenance. For example, the Senseless gene in Drosophila melanogaster and Pag3 in Caenorhabditis elegans are orthologous to Gfi1. Senseless and Pag3 are critical for neuronal development and sensory organ specification. Deletion of Gfi1 causes deficiency in inner ear hair cells in mice, giving rise to hearing loss as well as reduced numbers of cochlear neurons. Gfi1 also is expressed in cells of peripheral nervous system,as well as inner ear hair cells, the eye, rare pulmonary neuroendocrine cells (Kazanjian et al., 2006).

To fully understand how differential expression of genes that direct development of both lymphocytes and neuronal cells can produce very different cell types, analysis of mechanisms of functionally relevant regulatory genes should be determined. Evaluating shared and lineage specific mechanism may shed light on common and cell type specific control mechanisms in these distinct cell types. Genes that are controlled via shared regulatory mechanisms in different cell types might be major determinants of differentiation. A selection of regulatory genes which share transcriptional regulation by CTCF reveals both shared and distinct elements.

## METHODS

### Comparative ChIP-Seq analysis

Publicle available CTCF ChIP-Seq data files are extracted from Gene Expression Omnibus (GEO) database. GEO numbers of each data set are given in Table 1. Data is retrieved in UCSC Genome Browser (https://genome.ucsc.edu/) as custom tracks. UCSC gene references are used for gene annotations. Data is visualised according to mm9 assembly.

**Table 1.** List of publicly available CTCF ChIP-Seq data.

| Cells | GEO Accession Number |
| --- | --- |
| Neural progenitor cells | GSE65748 |
| Differentiated motor neurons | GSM1468394 |
| Pro B cells | GSM1023420 |
| Double positive T cells | GSM1023418 |
| Double negative T cells | GSM1023416 |

### Phylogenetic analysis

The phylogenetic trees for genes were extracted from the Ensembl database from the Comparative Genomics section using Gene Tree option.

Gene Tree IDs are given at Table 2 for genes under investigation:

**Table 2.**
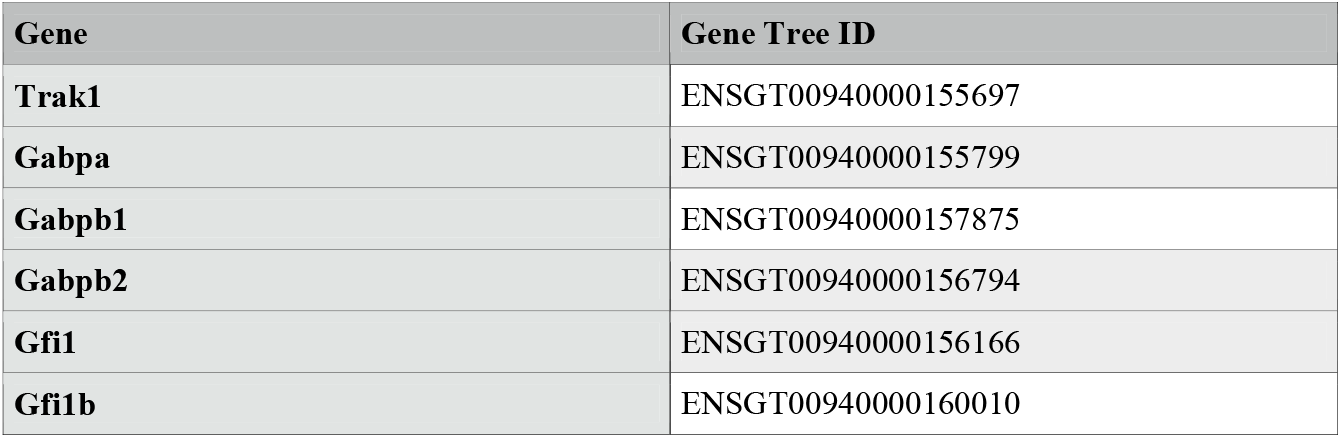
Gene Tree IDs.

### Comparative gene expression analysis

Gene expression levels are retrieved from the study GNF Mouse GeneAtlas V3 with GEO Accession number of GSE10246. GEO2R tool under NCBI GEO website was used to visualise gene expression levels of selected samples.

## RESULTS AND DISCUSSION

In this study, we performed epigenetic analysis of CTCF binding Trak1, Gabpa, Gabpb1, Gabpb2, Gfi1, Gfi1b gene loci at T and B lymphocytes at different developmental stages, as well as in neural progenitor cells and motor neurons. Genes of interest fell into two categories: a role in mitochondrial biology (Trak1, Gabpa, Gabpb1 and Gabpb2) or differentiation factors (Gfi1, Gfib). Mitochondria play critical roles in both neuronal and lymphocytic lineages, but the roles are different in the two lineages. Similarly, neuronal and lymphocytic lineages undergo multistep differentiation events leading to functionally diverse cell populations. Both lineages are affected by CTCF binding to genes that play a role in differentiation, so comparison of regulatory events in representative genes may begin to illuminate shared and distinct pathways.

We compared expression levels of these genes in brain and lymphocyte populations using publicly available data and correlated these results with differential CTCF binding. Finally, phylogenetic analysis was performed for genes under investigation to identify highly conserved motifs. In these analyses, we observed common CTCF binding events as well as differential binding of CTCF in various cell types at different stages. When differential binding events are compared to gene expression levels, expression levels correlated with selective binding or non-binding of CTCF at these genes. Genes under investigation in this study play critical roles in development of both lymphocytes and neurons. Significantly, some genes epigenetically regulated by CTCF exhibited similar control mechanisms during differentiation of neurons and lymphocytes, but these analyses also revealed use of lineage specific binding domains.

### Genes Implicated in Mitochondrial Biology

Trak1 is implicated in mitochondrial trafficking, which would be expected to be more important for neurons than lymphocytes. There are 17 CTCF binding sites at Trak1 gene loci and all 17 are occupied in double positive (DP) T lymphocytes, whereas only 13 binding sites are observed at double negative (DN) thymocytes which represent the prior stage of T cell differentiation (Figure 1). Therefore, CTCF plays a role in regulating T cell differentiation at this stage of differentiation. Elevated binding of CTCF at Trak1 gene from double negative stage to double positive stage might suggest Trak1 plays an important role in DP T cell differentiation. Given that the Trak1 gene is not expressed thymocytes and T cells, Trak1 cell expression appears to be suppressed in T cells, This absence of Trak1, which is thought to regulate mitochondrial trafficking, suggests that polarization and localization of mitochondria might not be critical in thymocyte selection, but it does play a role in B cells.

**Figure 1.**
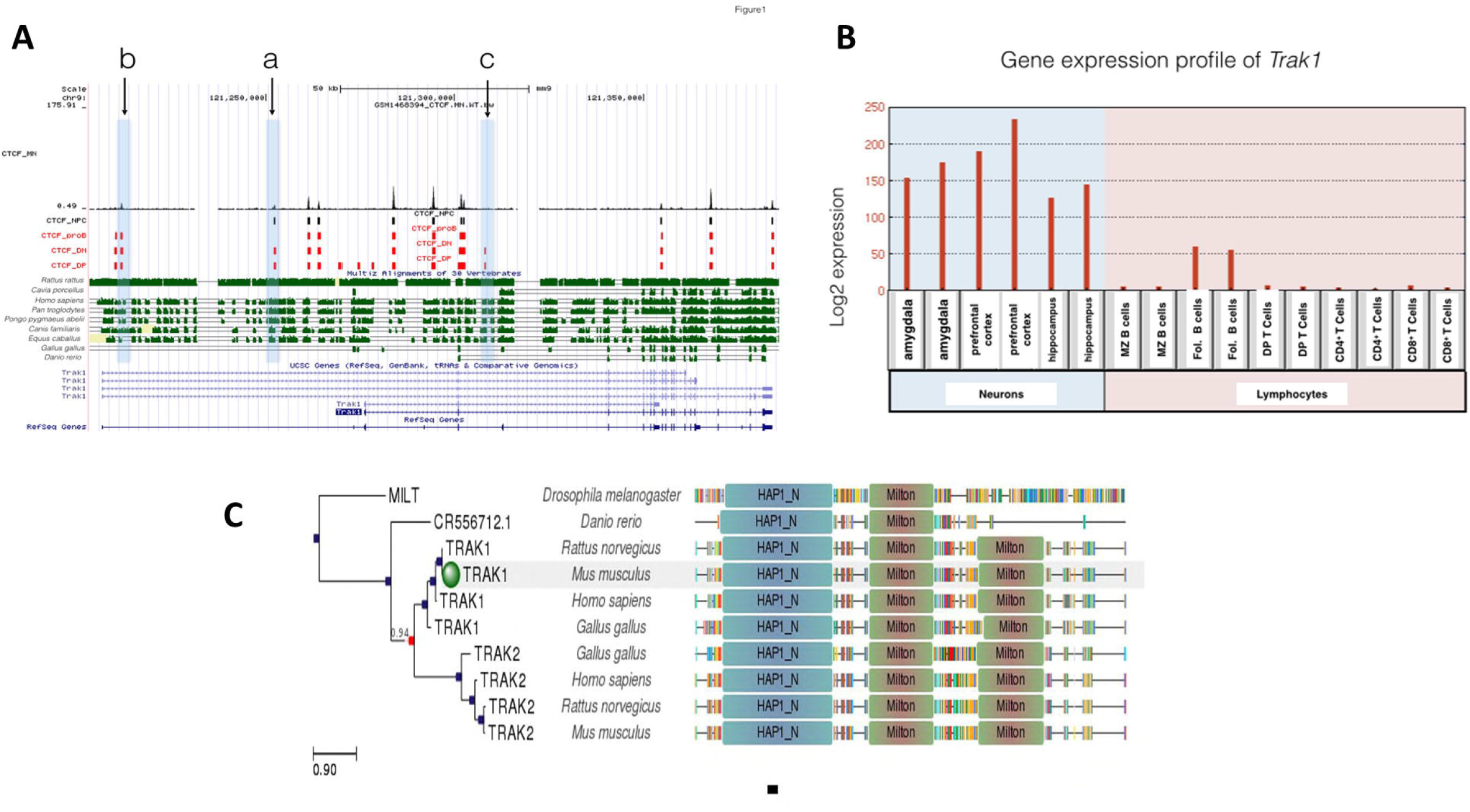
**A:** Gene conservation and CTCF binding profile at *Trak1* locus, **B:** *Trak1* gene expression in neurons and lymphocytes, **C:** *Trak1* gene phlogeny depicting conserved domains.

There are 10 different CTCF binding events at Trak1 gene in proB cells. All of these events are shared among DP and DN T cells. Although the binding profile differs between DP and DN T cells, those 10 shared binding events in proB and T cells suggest tight control of Trak1 gene expression during both B and T lymphocyte differentiation (Figure 1). However, there is a diference in gene expression profile, which reveals that Trak1 is expressed in follicular B cells but not in T cells. Those CTCF binding sites that are occupied in T cells, but not in B cells are likely to be important for suppression of Trak1 gene expression in T cells and suggest that polarization and localization of mitochondria is critical for B-cell differentiation. As sites ‘a’ and ‘c’ in Figure 1 are occupied in DN and DP T cells; but not in B cells, this suggests that they are permissive for expression in the B cell lineage. CTCF contrubutes lymphocyte differentiation, specifically V(D)J recombination that ensures lymphocyte variability (Seitan et. al., 2012). Therefore, presence of several sites with differential binding of CTCF in the Trak1 gene suggests that transcriptional regulation of this gene might be of importance in lymphocyte differentiation with suppression in T cell lineage that is absent in the B cell lineage.

Unlike the T-cell lineage, Trak1 expression is critical for neuronal development. The Trak1 gene is expressed at high levels in amygdala, hippocampus and prefrontal cortex, which is consistent with a requirement for regulation of mitochondrial trafficking. Whereas, 9 of the CTCF sites in the Trak1 gene are shared for all cell types under investigation, the gene is expressed only in neurons and in B cells, with the highest levels in the brain. Only site c in Figure 1, is specific to DN and DP T cells; and absent in B cells, neural progenitor cells and motor neurons that express Trak1. Although there 17 potential binding sites for CTCF in the Trak1 gene, binding at site c is the primary site for regulation of expression by CTCF. A lack of binding for CTCF at site c is necessary for expression, but binding of CTCF at site a may increase Trak1 expression in cells without CTCF at site c. In contrast, there is differential binding at sites a and b in Figure 1 in lymphocytes and neural progenitor cells. Binding is seen at site a for neural and T-cell lineages, but not in B cells. In contrast, both B and T cells as well as mature motor neurons exhibit binding at site b, but there is no signal at site b in neuronal progenitors. Due to limitations of publicly available data, gene expression levels only in mature neurons were investigated in this study. Thus, site c is a critical component in the expression of Trak1, with other sites or other transcription sites responsible for modulating the levels of expression. Common and shared CTCF binding events at Trak1 gene suggest further transcriptional regulatory factors that control Trak1 gene expression levels that differ in neurons and lymphocytes. Identification of factors that bind at site c in neurons, neural progenitors and B cells, may illuminate additional differentiation factors for Trak1.

This conclusion is reinforced by phylogenetic analysis. When we analyzed CTCF binding sequences at sites a, b and c of the Trak1 gene locus across species, we observed that site c is the most highly conserved sequence across species including *Rattus rattus, Homo sapiens, Pan troglodytes, Pongo pygmaeus abelii, Canis familiaris* and *Equus caballus*. Site a and b are also conserved, but conservation of site c is more prominent. This well-conserved site has specific CTCF binding in DN and DP T cells. Whereas, site a shows high similarity in species *Homo sapiens, Pan troglodytes* and *Pongo pygmaeus abelii*. These two sites have differential CTCF binding profiles in B lymphocytes and neural progenitor cells. However, these three CTCF binding sites are not conserved in *Gallus gallus* and *Danio rerio* species, which suggests that this aspect appeared at the mammalian level and is absent in organisms whose immune system is regulated differently.

Unlike Trak1, Gabpa has only two CTCF binding sites (Figure 2A). Given the role of the GABP transcription factor in mitochondrial biogenesis (Yang et al, 2014) and its expression in all the lineages examined (Figure 2B), the limited extent of CTCF binding events was not expected. The CTCF binding sites are occupied for T-lymphocyte differentiation, but not for neuronal or B-cell lineages. In particular, the Gabpa gene exhibited a specific CTCF binding site in DP T cells shown in Figure 2A at site a and a specific binding event at site b in DN T cells. Transcriptional regulation of this gene might be of importance in T lymphocyte differentiation, given its higher expression levels in the T-cell lineage. However, there were no binding events observed in motor neurons, neural progenitor cells and pro B cells. No differences are observed between Gabpa gene expression levels in DP and CD4+ and CD8+ T cells, although expression levels were generally higher in T-cell lineages than in B-cell and neuronal lineages. Gabpa gene expression in neurons and lymphocytes appear to be regulated independent of CTCF, which is consistent with limited conservation of CTCF binding sites in phylogenetic studies. Site b is only conserved between human and Equus caballus. Whereas, site a is well conserved among Homo sapiens, Pan troglodytes, Pongo pygmaeus abelii and Equus caballus (Figure 2C). Although our phylogenetic analysis revealed that Gabpa gene has the highest overall conservation profile among all genes under investigation, the absence of conserved CTCF binding is consistent with suggestions that Gabpa gene expression is regulated independent of CTCF.

**Figure 2.**
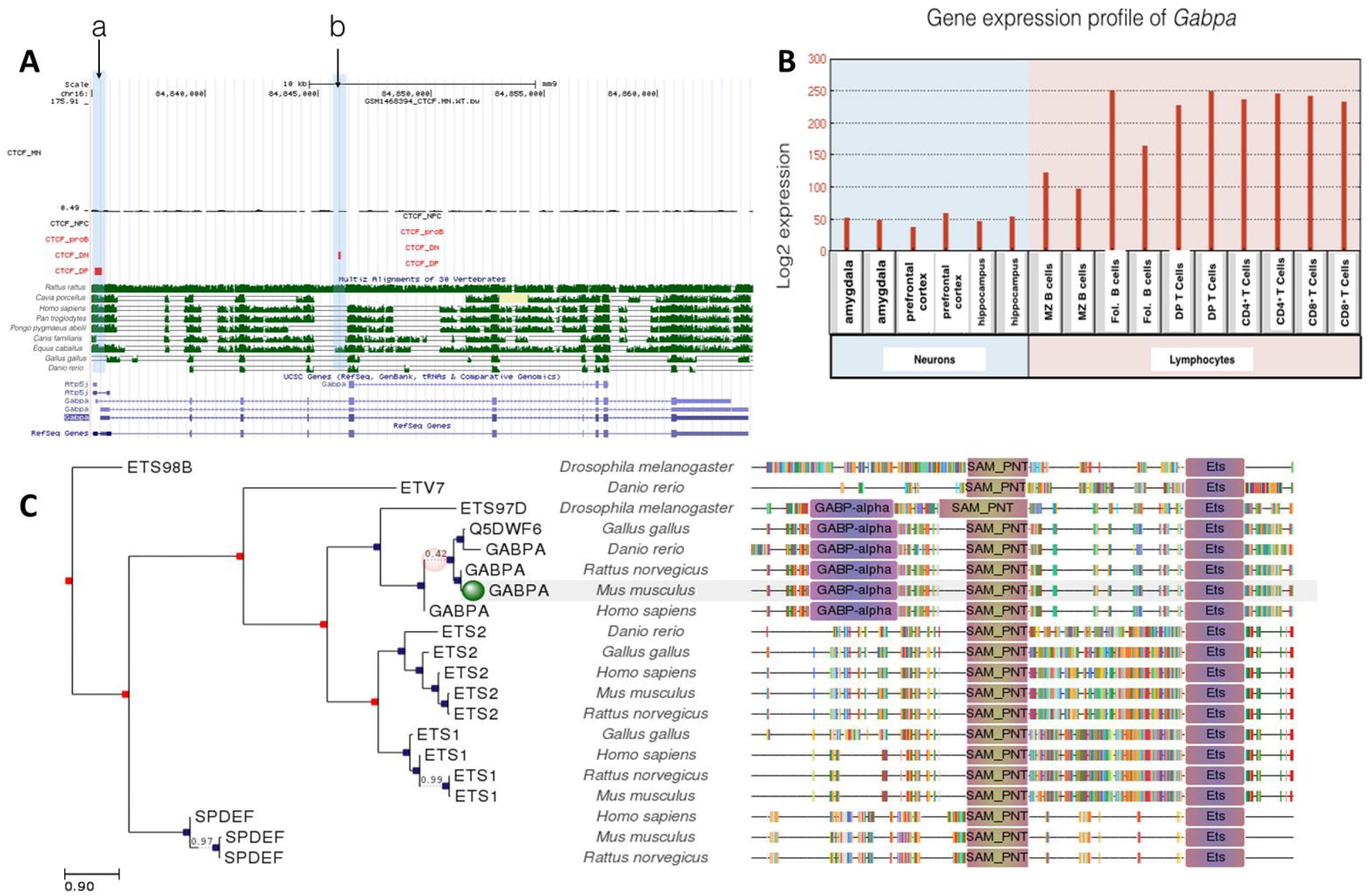
**A:** Gene conservation and CTCF binding profile at *Gabpa* locus, **B:** *Gabpa* gene expression in neurons and lymphocytes, **C:** *Gabpa* gene phlogeny depicting conserved domains,

In contrast to Gabpa, Gabpb1 gene includes a common CTCF binding site shared with neurons and lymphocytes. Site a in Figure 3A is occupied in all cell types except proB cells, while site b is occupied in all cell types examined. Interestingly, Gabpb1 gene expression is 2 to 3-fold higher in follicular B cells than thymocytes and expression in neurons is lower still (Figure 3B). CTCF binding at site ‘a’ might regulate suppression of Gabp1 in T cells and neurons along with other transcription factors. Correlation of CTCF binding analysis and gene expression profile suggests that CTCF bindings play a role in epigenetic regulation of Gabpb1 gene.

**Figure 3.**
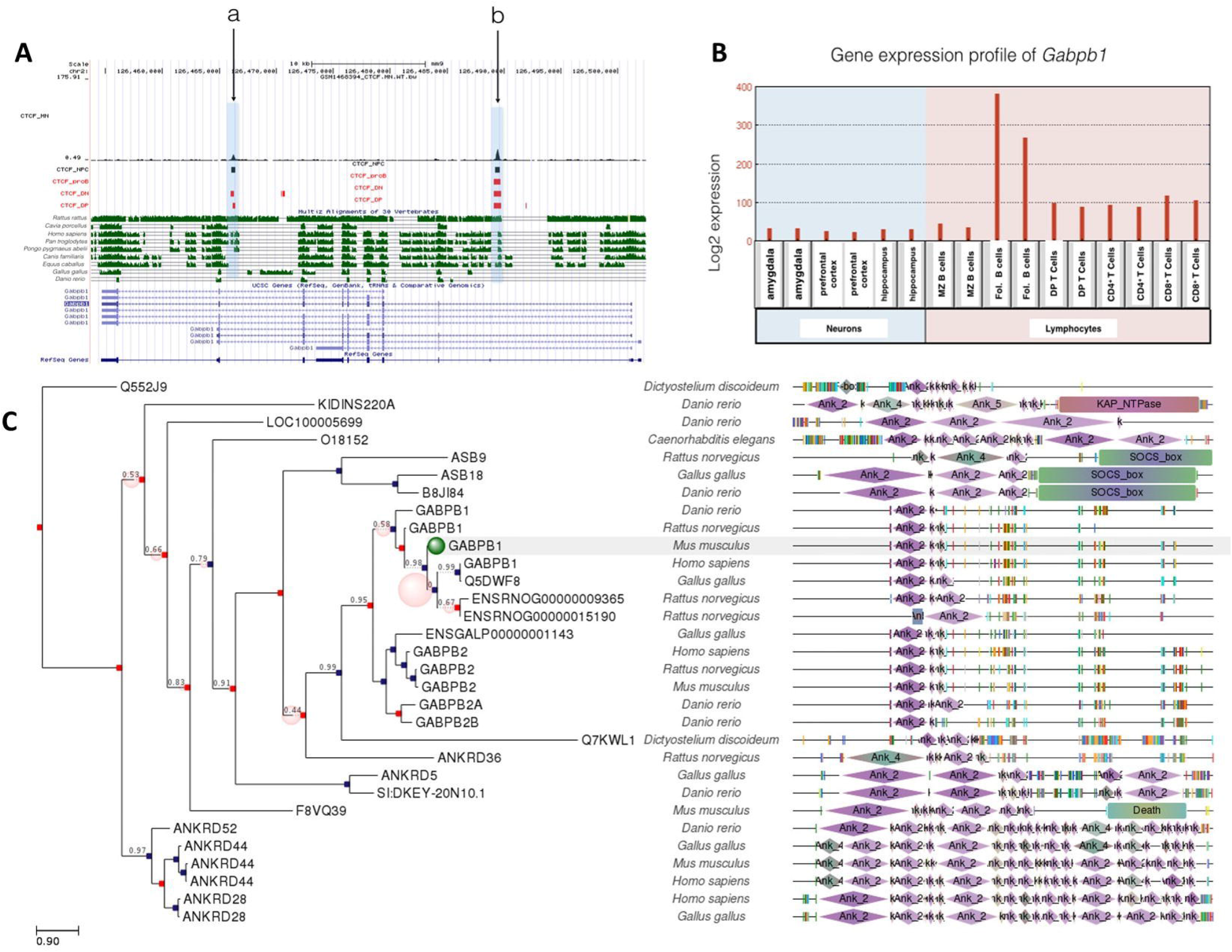
**A:** Gene conservation and CTCF binding profile at *Gabpb1* locus, **B:** *Gabpb1* gene expression in neurons and lymphocytes, **C:** *Gabpb1* gene phlogeny depicting conserved domains,

A common CTCF binding event (site b) at Gabpb1 gene is well conserved in Rattus rattus, Homo sapiens, Pan troglodytes, Pongo pygmaeus abelii, Canis familiaris and Equus caballus. Site a, which is occupied in neurons and T cells; is conserved among Rattus rattus, Homo sapiens, Pan troglodytes, Pongo pygmaeus abelii.

Gabpb2 gene includes a CTCF site common to all cells examined. There is only one CTCF site ‘site a’ which is only occupied in DP T cells. Like Gabpa and Gabpb1, Gabpb2 is differentially expressed in neurons and lymphocytes (Figure 4). Gabpb2 is highly expressed especially in B lymphocytes and moderately expressed in T lymphocytes, but has almost no expression in amygdala, hippocampus and prefrontal cortex. Similar to Gabpb1, Gabpb2 expression is also suppressed in neurons than lymphocytes. However, CTCF binding site at Gabpb2 is conserved among more species than Gabpb1. This site is well conserved among *Rattus rattus, Cavia porcellus, Homo sapiens, Pan troglodytes, Pongo pygmaeus abelii, Canis familiaris* and *Equus caballus*. The role of CTCF binding in regulating Gabpb2 in these cells is modest and may require interactions with other transcription factors.

**Figure 4.**
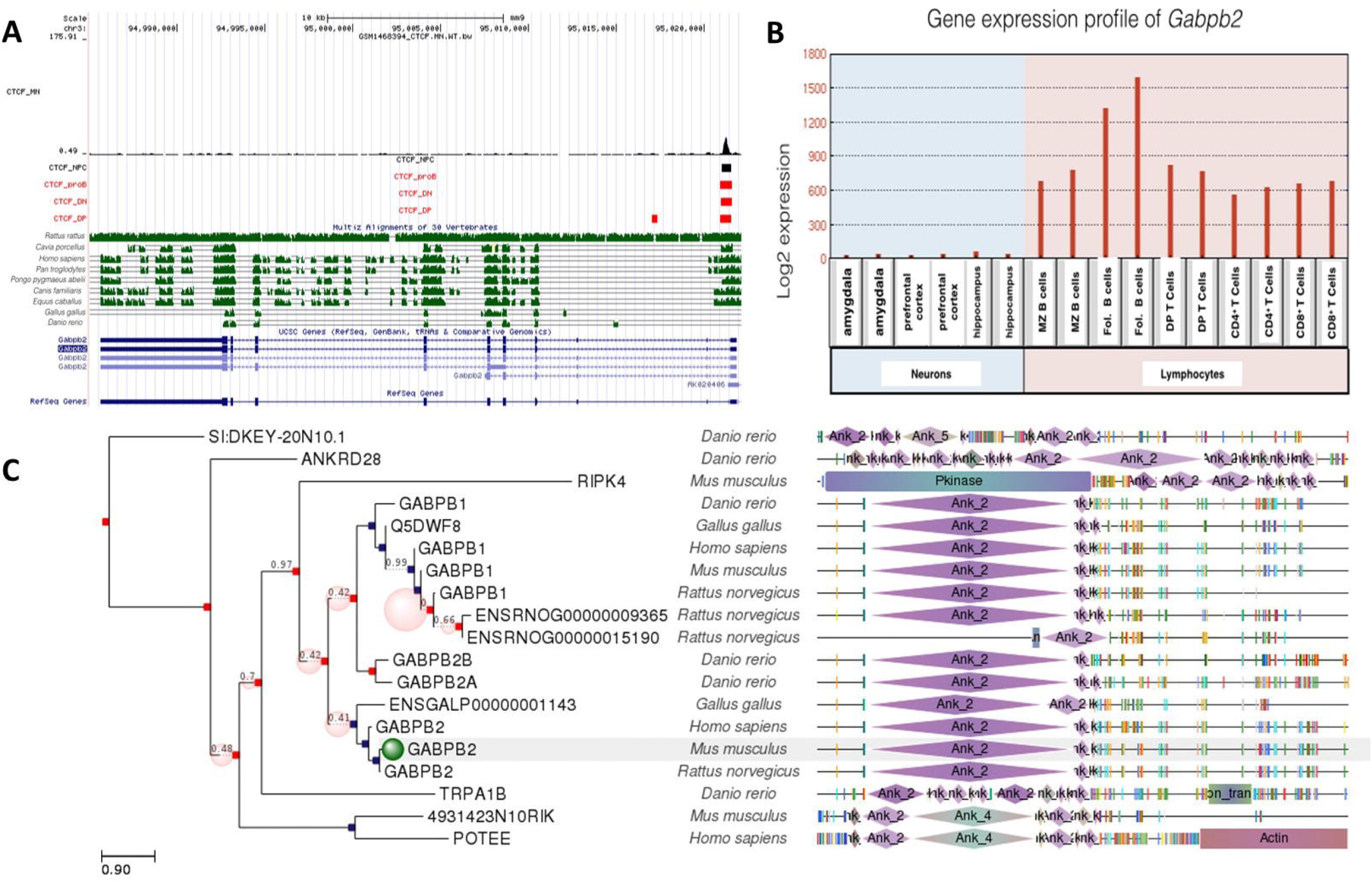
**A:** Gene conservation and CTCF binding profile at *Gabpb2* locus, **B:** *Gabpb2* gene expression in neurons and lymphocytes, **C:** *Gabpb2* gene phlogeny depicting conserved domains.

CTCF binding sites at Gabp2 are evolutionarily more conserved than those at Gabp1, which may indicate that Gabpb1 has more specific roles in mediating neurogenesis and lymphocyte differentiation. Moreover, higher conservation of gene regulation via CTCF in primates suggest that transcriptional suppression of Gabpb1 gene might contribute to neuronal diversity in primates.

Gfi1 and Gfi1b are zinc-finger transcription factors that play a critical role in hematopoiesis (cite_van der Meer et al 2010), but Gfi1b is also expressed in neurons, while Gfi1a is low or absent from neurons (Figure 5 and 6). Both DP T lymphocytes and proB lymphocytes have occupied by CTCF. There are 3 CTCF binding sites at Gfi1a gene locus for DP T lymphocytes, but only one is shared with pro B lymphocytes. Gfi1a gene expression is relatively higher in DP T lymphocytes than other lymphocytes and neurons. This might be due to several factors. Given the fact that CTCF plays critical roles in differentiation of lymphocytes, elevation of gene expression levels might be dependent of CTCF binding to these sites.

**Figure 5.**
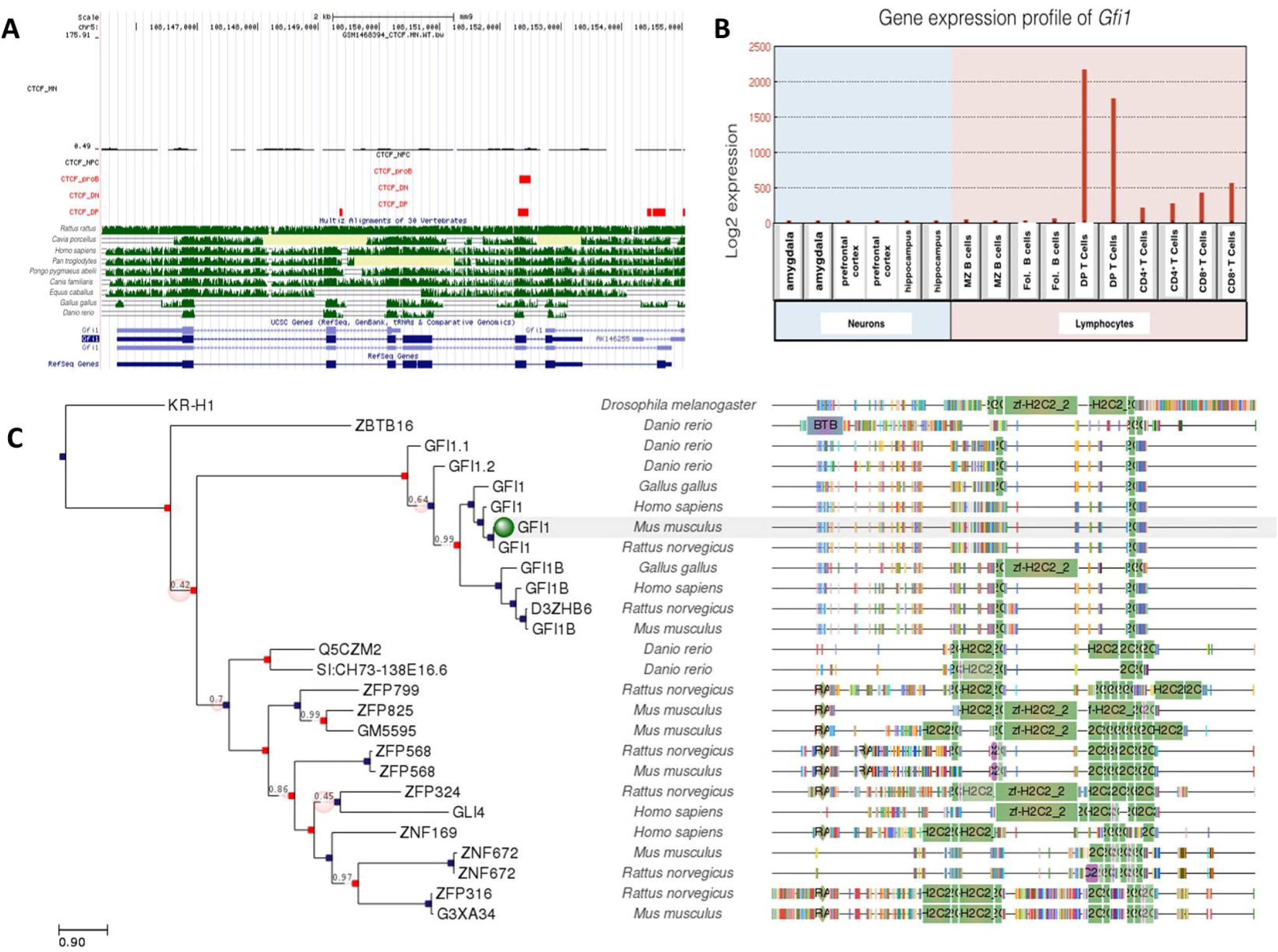
**A:** Gene conservation and CTCF binding profile at *Gfi1* locus, **B:** *Gfi1* gene expression in neurons and lymphocytes, **C:** *Gfi1* gene phlogeny depicting conserved domains,

**Figure 6.**
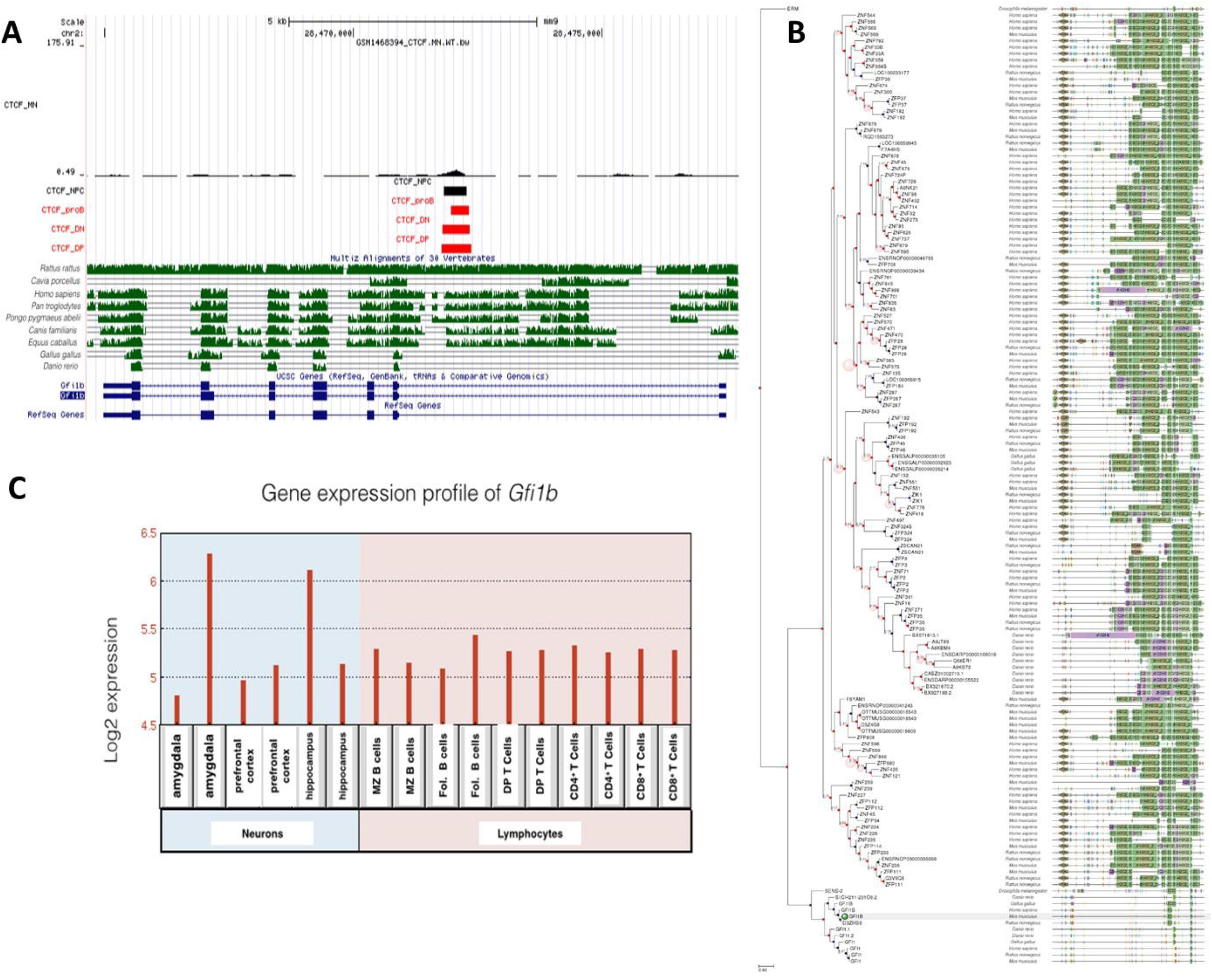
**A:** Gene conservation and CTCF binding profile at *Gfi1b* locus, **B:** *Gfi1b* gene phlogeny depicting conserved domains, **C:**. *Gfi1b* gene expression in neurons and lymphocytes.

Unlike Gfi1, Gfi1b is expressed at moderate levels in both neuronal and lymphocyte lineages. Interestingly, there is only one conserved CTCF site at the Gfi1b gene locus, which is occupied in all cell types examined. Consistent with this, gene expression levels for Gfi1b are similar but not high in all cells. Both CTCF binding profile an gene expression levels are different than Gfi1. This might suggest that Gfi1 critical role in DP T cell differentiation, while Gfi1b is more important or neuronal differentiation as well as the lymphocytic lineage.

CTCF binding event at Gfi1 gene that is shared in B and T cells is conserved among *Rattus rattus, Homo sapiens, Pan troglodytes, Pongo pygmaeus abelii, Canis familiaris* and *Equus caballus* as well as *Gallus gallus* and *Danio rerio*. A more in-depth analysis of these conserved sites in *Gallus gallus* and *Danio rerio* will shed light in understanding of potential roles of Gfi1 in evolutionary variation of lymphocytes.

Gfi1a gene is phylogenetically well-conserved in many species. There are no CTCF binding occupied in neurons and neural progenitor cells. Low expression levels of Gfi1s in neurons indicate that regulation of this gene is CTCF-independent. Although Gfi1b is highly homologous to Gfi1, differences in expression levels suggest that Gfi1b is not only critical for lymphogenesis but also neurogenesis. The common CTCF site at Gfi1b gene is well conserved among *Rattus rattus, Homo sapiens, Pan troglodytes, Pongo pygmaeus abelii, Canis familiaris* and *Equus caballus*. Gfi1a domains are conserved in more species than Gfi1b, despite their homology, so it is considered to be the more ancestral gene.

Consequently, differences in the regulation mechanisms of neurogenesis and lymphopoiesis may contribute to the identification of new target and signaling pathways for neurodevelopmental diseases. In this case, there appear to be more parallels in transcriptional regulatory pathways between neuronal and B cell lineages than between T cell and neuronal lineages. This may reflect the common role of neurons and B cells as secretory cells. Regardless, this emphasizes the importance of selecting the appropriate targets and cell types in looking for plasma markers of neurodevelopmental and neurodegenerative diseases. In this context, specific transcriptional regulators that play a role in their different expression levels need to be identified, although they contain common CTCF binding sites in neurons and lymphocytes.

## Abbreviations

CTCF: CCCTC-binding factor
ChIP-Seq: Chromatin immunoprecipitation coupled with deep sequencing

## ACKNOWLEDGEMENTS

The authors have no conflict of interest.

